# Mixed Effects Machine Learning Models for Colon Cancer Metastasis Prediction using Spatially Localized Immuno-Oncology Markers

**DOI:** 10.1101/2021.08.01.454649

**Authors:** Joshua J. Levy, Carly A. Bobak, Mustafa Nasir-Moin, Eren M. Veziroglu, Scott M. Palisoul, Rachael E. Barney, Lucas A. Salas, Brock C. Christensen, Gregory J. Tsongalis, Louis J. Vaickus

## Abstract

Spatially resolved characterization of the transcriptome and proteome promises to provide further clarity on cancer pathogenesis and etiology, which may inform future clinical practice through classifier development for clinical outcomes. However, batch effects may potentially obscure the ability of machine learning methods to derive complex associations within spatial omics data. Profiling thirty-five stage three colon cancer patients using the GeoMX Digital Spatial Profiler, we found that mixed-effects machine learning (MEML) methods^†^ may provide utility for overcoming significant batch effects to communicate key and complex disease associations from spatial information. These results point to further exploration and application of MEML methods within the spatial omics algorithm development life cycle for clinical deployment.

## 1. Introduction

Practicing pathologists routinely order immunohistochemical stains to assess the spatial distribution of important biomarkers for disease prognostication. Recently, higher resolution spatially resolved technologies have emerged. Nature Methods declared “Spatially Resolved Transcriptomics” technologies as the method of the year in 2020 ^1^. Although multiplexed immunohistochemistry and laser capture microdissection approaches once served as popular methods for assessing spatially resolved omics information, these approaches are largely intractable for high throughput analysis because quantitative analysis is hindered by significant tissue distortion from chemical restaining processes, insufficient multiplexing, and prohibitive costs. New spatial omics technologies can multiplex far more markers through targeted cleavage of oligonucleotide tags ^2–8^. For instance, the Nanostring GeoMX Digital Spatial Profiler (DSP) first utilizes multiple immunofluorescent (IF) antibody stains to highlight various cellular lineages, then uses UV-light to cleave attached oligos for quantification via the nCounter instrument or next-generation sequencing (NGS)^9,10^. A particular novelty of this technology is the ability to highlight regions of interest (ROI) through segmentation of the IF stains or selection of specific locations of interest. Informatics techniques developed for spatial data include estimation of: 1) which genes are spatially variable, 2) how spatial clustering patterns of expression may differ between cases and controls, 3) co-localization and coordination between cellular populations, accomplished using canonical protein markers or cellular deconvolution, and 4) integration with other imaging (e.g., morphology, architecture via hematoxylin and eosin stain, H&E) and single-cell multi-omics (e.g., RNASeq, ATACSeq; models of cell fate) which can further shed light on diagnosis and prognostication ^11^.

Prospective deployment of clinical decision aids can use low-cost tissue staining techniques (e.g., IHC) to profile protein markers deemed relevant through spatial omics analysis. However, existing approaches (e.g., differential expression and spatial autocorrelation) suffer from technical factors, including: reagent application, batch effects, and other forms of repeated measurements. These confounders have the potential to bias disease associations and preclude prospective classifier deployment. Compared to traditional penalized, generalized linear modeling approaches, algorithms that can use the complete multiplexed set of markers and their interactions (e.g., Random Forest models) may be best positioned to capture the disease pathogenesis yet are likely less resistant to batch effects. Thus, we need machine learning algorithms that can fully utilize spatial multiplexed information while being immune to batch effect corruption. In this study, we assess the potential for mixed-effects machine learning (MEML) methods to address this issue, capturing unintuitive spatial and multiplexed complexities, while respecting patient and batch level differences.

We assessed colorectal adenocarcinoma (CRC) cases, with or without nodal and/or distant metastasis using spatial omics technologies (**Figure 1A-B**). Prognosis / metastatic potential is well characterized based on presence of tumor invasion into distinct tissue layers (e.g., epithelium, lamina propria, submucosa, *muscularis propria*, pericolic fat, and serosa). Previous research has demonstrated the importance of specific somatic alterations (e.g., APC, MMR) and tumor-infiltrating lymphocytes (and their spatial coordination) as prognostic indicators ^12–15^. Here, we used the DSP to collect spatial expression of 39 immune markers across 840 ROIs and applied MEML to characterize factors relating to different macro-architectural contexts (e.g., markers related to tissue interface with tumor, *inter*; within tumor, *intra*; away from tumor, *away*) and metastasis. This study will serve as the backbone of an expanded investigation of spatial immune markers descriptive of nodal and distant metastasis for Colon cancer patients.

**Figure 1:**
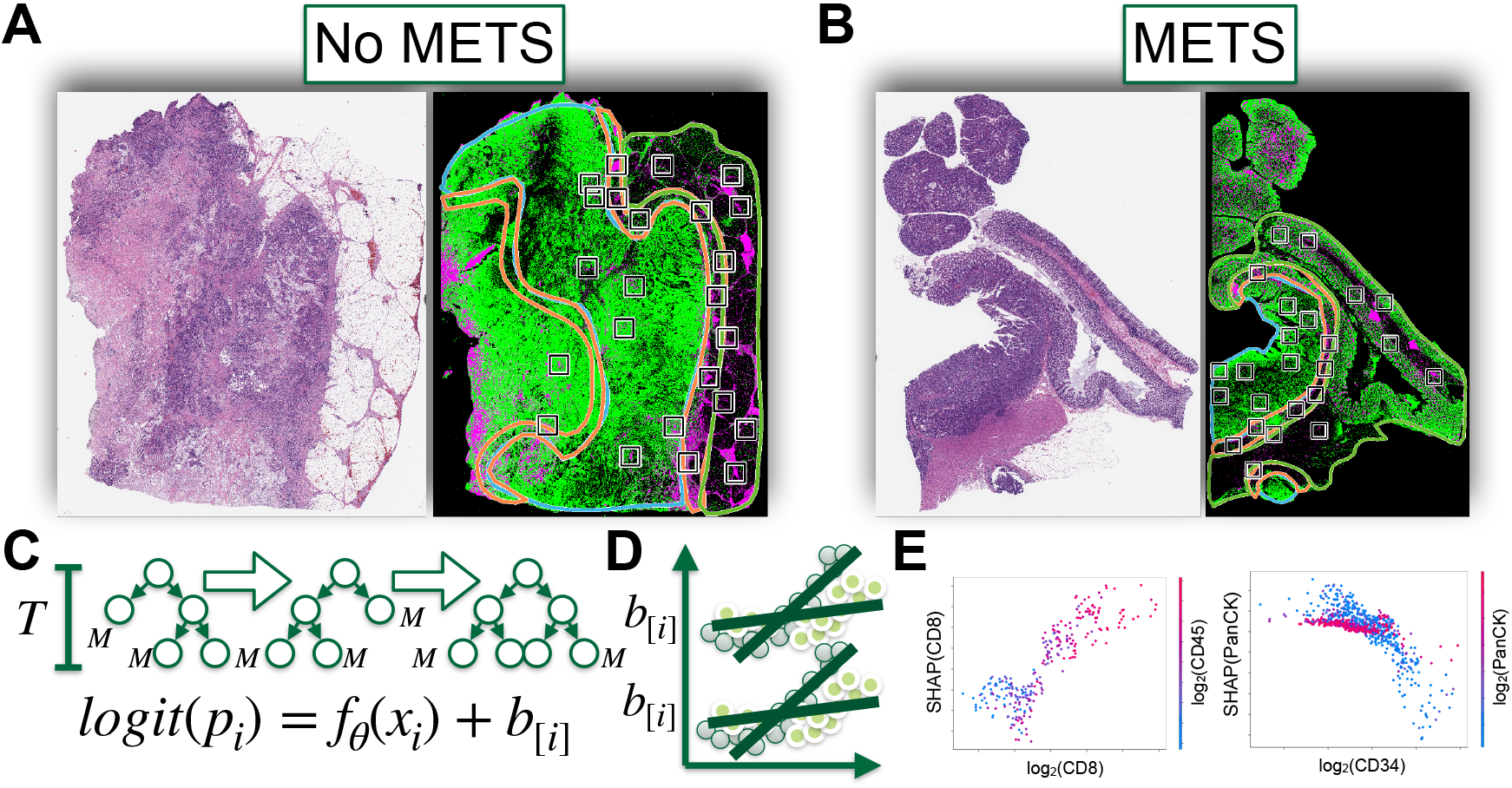
Overview of Experiment: Paired H&E and IF stains (PanCK stain, green; CD45 stain, pink) used to help select ROI (black squares) contained in macro-architectural contexts (outlined: blue for intra-tumoral, red for tumor interface, green for away from tumor) for: A) Non-metastatic patient, B) Metastatic patient; C) Tree boosting methods *f*_θ_(*x*_*i*_) combined with mixed effects modeling to adjust for D) patient/batch-level effects *b*_[*i*]_ (e.g., interactions within nested observations, continuous scale) to yield E) disease associated interactions; SHAP dependence plots demonstrate how predictor (x-axis) covaries with another (colors) and impact on predictor importance, y-axis

## 2. Motivation for Comparison Study

First, we will provide a broad overview of common analytical techniques for spatial omics data, then discuss potential statistical oversights (e.g., modeling of statistical interactions and nonlinearities, batch effects/repeat measurements), to highlight the advantages of MEML methods.

### 2.1. Review of Prior Spatial Omics Analysis Methods

Within a slide, there is spatial variation in marker abundance, such that regions of interest (ROI) that are closer together may exhibit similar gene/protein expression ^5,8,16^. Spatial autocorrelation is typically assessed using a Gaussian Process (GP), a mixed-effects model with a parameterized kernel. Included is an example of this model for multivariate normally distributed expression data:

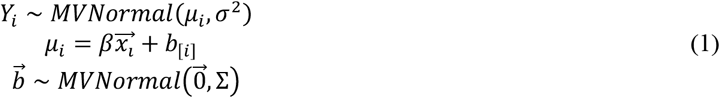

Where 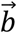 is a vector of random intercepts, here observed at locations 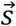. The covariance matrix of the random effects, Σ, is a kernel used to test spatial hypotheses; for instance, that the correlation in expression between two spatially dependent markers drops off with distance *d*, Σ_*i,j*_ = *a*^2^*exp*(− *ρ*^2^*d*(*s*_*i*_, *s*_*j*_)^2^). The target of inference would be *ρ*, the degree to which the correlation in expression is retained as a function of the distance (*d*; e.g., L2-norm) between the ROIs *i* and *j*. Less parametric forms of this hypothesis can be tested through network autocorrelation and, more recently, through prediction models such as graph neural networks (GNN), where the latter is useful for increasing the resolution of the spatial expression patterns– e.g., predicting the current ROI’s expression from that of neighboring (neighborhood *N*) cells/ROIs, *j* ^17,18^: *Y*_i_ = σ(Σ_*j∈N*_ *a*_*j*_*Y*_*j*_).

Other analyses expand the aforementioned GP models to estimate characteristic clustering patterns, enabling inter-slide comparison. Less parametric forms that take into account regional dependencies include graph neural networks, which can utilize aggregation/pooling mechanisms across nested observations to predict a single outcome while learning representative patterns. Here, establishing differential expression for specific markers and applying penalized regression procedures across a set of markers are simpler analytic approaches that compare spatially variable markers across slides and differentiate disease status. The latter is formalized below, where *Y*_i_ represents whether the case underwent metastasis, predicted from markers 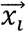:

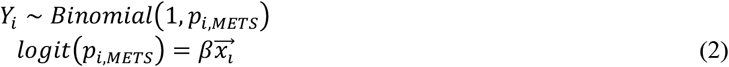

In this simplified scenario, spatial clustering patterns can be incorporated into this analysis by estimating stratified or interaction effects by assigned macro-architecture (e.g., *intra, inter, away*).

### 2.2. Motivation for Mixed Effects Machine Learning Approaches

The goal of this preliminary comparative study is to explore methods for classification of tissue macro-architecture/metastasis from spatially resolved expression patterns and to discuss opportunities for classifier development. In this section, we discuss deficiencies of existing methods. Machine learning (ML) techniques search for the ideal specification of interactions and nonlinear transformations that capture the relationship between the input and target of inference. For instance, classification and regression models (CART; e.g., Random Forest) utilize conditional decision splits to extract these associations ^19^. Interpretation methods (e.g., Shapley Additive Feature Explanations, SHAP) can derive disease-associated predictors and interactions ^20^. As omics datasets are usually high-dimensional and multi-collinear, CART methods excel in spotting relevant interactions.

However, these algorithms assume that observations are independent and identically distributed (*i.i.d.*), which is invalid in the presence of repeated measurements, where statistical dependencies exist between observations. In the case of spatial omics modalities, an ROI may exist within several levels of nested dependencies (e.g., ROIs per section, sections per block and/or profiling batch). Multiple blocks may be extracted from different biopsies across space (sampling site) and time. These factors may potentially exacerbate the negative impact of technical effects on statistical inference. Even after documenting and controlling for batch effects, the degradation in performance resulting from the correlation between batch and outcome has not been thoroughly studied in this context ^21,22^. Hierarchical modeling can leverage information across clusters of observations for efficient modeling of the fixed effects while respecting the dependency structure to avoid potentially biased inference. A random-intercept model, modeling different baseline measurements per cluster, follows the form of a linear mixed-effects model, which is a special case of the GP model ^23^:

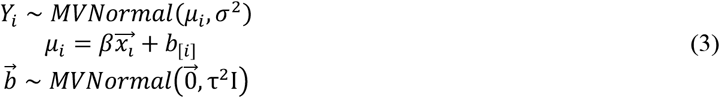

Given that spatial omics data may require algorithms that leverage nonlinear, complex high-order interactions offered by machine learning while respecting the dependency structure of the data, practitioners of spatial omics analyses, specifically those developing classifiers, should consider machine learning models which utilize mixed-effects modeling. Several such methods have been developed in the previous decade, being usually formulated as ^16,24–26^:

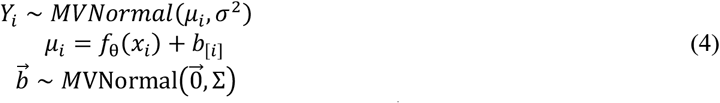

This modeling approach learns to estimate a random-effect component, 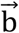, that allows the estimation of a fixed-effect component robust to clustering by replacing a linear fixed-effects component, *βx*_*i*_, with the fixed-effects machine learning model, *f*_θ_(*x*_*i*_) (**Figure 1C-D**). Model fitting is typically accomplished by first removing the random effects from the outcome before fitting the fixed effects machine learning model, 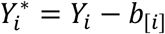, and then estimating or drawing fixed effects components through frequentist or Bayesian procedures. This procedure is repeated until convergence of both the fixed and random effects components. Potential drawbacks of separating the fixed and random effect components include having to omit the random-effect component during inference, where in many cases out of cluster prediction yields similar results to the fixed effects counterpart, and complexities in the random effects are not well captured (only random intercepts and slopes). Alternatively, clustering can be directly included in the model decision splits. For instance, for a given split, one can fit a generalized estimating equation model with a custom dependency structure to assess the ability to partition variable *k* with cutpoint 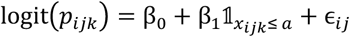, where ϵ_*ij*_ ∈ *N*(0, Σ) and the correspondent Wald statistic on *β*_1_ may inform the variable and value to split on ^27^.

Failing to consider the dependency structure of omics data for ML risks introducing bias by learning properties of the batch rather than the inference target. As omics data becomes increasingly available at the single-cell and spatial resolutions, mixed-effect modeling approaches may be optimally positioned to capture potential dependency structures/repeat sampling in the data.

## 3. Materials and Methods

### 3.1. Data Acquisition and Preprocessing

Under IRB approval, pathology reports and slides were reviewed for colon adenocarcinoma cases at Dartmouth Hitchcock Medical Center (DHMC) from 2016 to 2019. A total of 36 biopsies were initially selected for spatial profiling, half of them exhibiting local invasion but no nodal or distant metastasis (no METS) and the other half of them having undergone nodal and/or distant metastasis (METS). The cohort was restricted to stage 3 assignments under the pTNM staging system, which balances the impacts of local invasion, nodal and distant metastasis for prognostication. Cases were matched between the non-metastatic and metastatic groups based on tissue size (measured through connected component analysis of whole slide image), tumor grade, mismatch repair (MMR) dysregulation status, site of the tumor (e.g., left or right colon), age, and sex. Matching was achieved through conducting fisher’s exact tests and two-sample t-tests after iterative resampling. For instance, average ages for the no METS and METS groups were 71±13 and 66±16 respectively (p=0.34) and stratified by sex, average ages for females and males were 68±18 and 69±13 respectively (p=0.96). A similar percentage of males comprised the METS and no METS groups (56% and 58% respectively, p=1.0). Tissue blocks were sectioned into 5-μm sections. Sections were stained with fluorescent-labeled antibodies (highlighting tumor (PanCK), immune cells (CD45), and nuclei (SYTO13)) which are covalently linked to photocleavable oligonucleotide tags targeting a panel of Immune Cell Profiling and Tumor Immune Environment markers. These sections were visualized using immunofluorescent (IF) images provided by the digital spatial profiling (DSP) instrument. Subsequent sections were stained with hematoxylin and eosin (H&E) and scanned using the Leica Aperio-AT2 scanner at 20x. IF whole slide images (WSI) were stored in TIFF format (16-bit unsigned color channels; one channel per stain) and H&E images were stored in SVS format (8-bit color channels). IF and H&E images were viewed simultaneously by the pathologist using the ASAP annotation software, which allowed the pathologist to delineate immune populations within macroarchitectural regions of interest (*intra* or within tumor, *inter* or at interface with tumor, *away* from tumor) on the IF images using polygonal/spline annotations. For each slide, a semi-automated workflow placed eight regions of interest (ROI; square grids of maximal spatial dimensions allowable by the GeoMx DSP instrument) within each annotation region (24 ROI per slide) after removing ROI with low immune cell density as quantified by the presence of CD45 stain. ROIs were mapped onto IF slides, uploaded to the DSP, and manually registered to the unmanipulated DSP IF images. ROI were placed onto DSP slides corresponding to the semi-autonomous placements and manually adjusted to the nearest appropriate region by a practicing pathologist if any ROI were misplaced (e.g., incorrect region or low immune cell density). Within each ROI, immune cells were segmented by using color thresholding and connected component analysis to deduce which areas in the ROI had overlapping nucleus and immune stains while excluding the tumor stain. The oligo tags within the immune cell areas were cleaved using ultraviolet light, collected and counted with nCounter to quantify protein expression within the immune cell regions of all ROI. Four slides were assessed at a time due to the DSP’s batching mechanism. Additional cases were included/excluded due to tissue lifting after cover-slipping procedures (leading to case removal) or to balance subsequent batches, leading to inclusion of 35 cases (840 ROIs).

A total of 28 ROIs were removed after quality control procedures, retaining 812 ROIs. Within each batch, we separately performed and compared expression after ERCC normalization, normalization based on nuclei area and count, and normalization to housekeepers and IgG isotype controls. Using the raw data, we compared log2-transformed counts of housekeeper markers to each other (S6, Histone H3, GAPDH) and separately compared counts of IgG isotope controls (Rb IgG, Ms IgG1, Ms IgG2a), also looking for significant batch effects. We used this information to sub-select markers that were consistent with other control markers and exhibited the lowest potential for batch effects (Housekeepers: GAPDH, Histone H3; IgG: Rb-IgG, Ms-IgG1), and normalized the expression of the other markers based on the geometric mean of these respective markers. A preliminary analysis demonstrated the lowest impact from technical factors when using IgG isotopes, so we normalized expression based on IgG isotope controls for the following experiments. ROIs were further labeled with positional coordinates, nuclei count and total area, MMR alteration status, age and sex, site of origin/metastasis, tumor grade, nodal and distant metastasis status, and macro-architectural region (*intra*, *inter*, *away*). A total of 36 protein markers were selected for analysis, not including housekeeping genes. We emphasize that the collected data represents a preliminary analysis and does not represent the data for final classifier development. Nonetheless, the data collected may prove helpful for the comparison of different classification algorithms.

### 3.2. Experimental Design: Prediction Tasks and Modeling Approaches

We compared predictive performance of several mixed-effects machine learning (MEML), two fixed effects techniques (Random Forest, XGBoost) ^28^, and classical statistical methods. As input to the algorithms are 36 markers (ROI-level fixed effects), age and sex (patient-level fixed effects), batch (random effects), and ROI coordinates (kernel). First, we treated the patient as a random intercept and predicted whether ROI could be localized to the tumor interface (*macro*), estimating: a) out of cluster predictive accuracy (*macro-ooc*; tested on held out set of patients) and b) within batch accuracy (*macro-ws*; samples from the same batch were partitioned for testing). Then, using the profiling batch as a random intercept, we sought to establish whether, based on assigned macro-architectural region, colon cancer metastasis could be predicted (*mets*); first, predicting *mets* regardless of assigned macro-architecture (*mets-overall*; including whether ROI was intra- or inter-tumoral as additional predictors). We then built classifiers to predict whether there was nodal or distant metastasis based on localization of spatial markers within distinct macro-architecture (*mets-intra, mets-inter, mets-away*). Our primary comparison was to compare out-of-cluster sampling performance with within-cluster sampling for the macro tasks. Since predicting out-of-cluster leads to the exclusion of random effects, we expect differences between fixed effects machine learning models and their mixed effects counterparts to be marginal. Thus, for the *mets* tasks, we focused on comparison of predictions within batch, as we wanted to shift focus from development of a classifier which can be used on external datasets to one where interrogation of important fixed effects predictors could potentially yield metastasis-related factors worth follow up. We provide brief overviews of selected modeling approaches and their implementations (R v3.6, Python v3.7):

1. **Gaussian Process Boosters (GPBoost):** tree boosting algorithms are additive models which fit weak learners to the residual between the outcome and sum of previous trees. Covariance parameters that account for random intercepts and arbitrary covariance functionals are updated at each boosting step via iterative minimization of the model likelihood and risk functional by gradient/newton descent. GPBoost models can generalize to classification tasks by regressing on a continuous latent outcome, dichotomized to yield binary outcomes. We model patients/batches as grouped random intercepts ^24^:

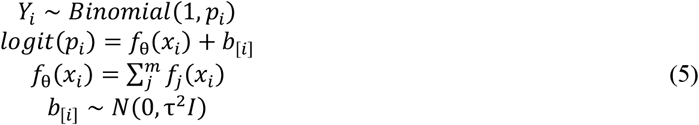
2. **GPBoost with Nested Coordinates (GPBoost-Coords):** Similar to GPBoost, with a random intercept for patient/tissue section and grouped within-patient, a spatial Gaussian Process with covariance function of exponential form which captures spatial dependence between ROI:

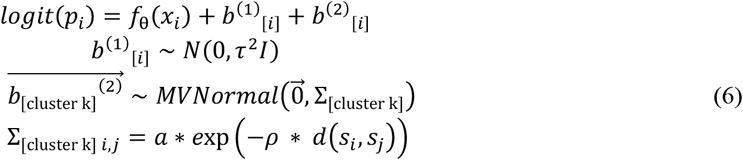
3. **Semi-Parametric Mixed Effects Bayesian Additive Regression Trees (SP-BART):** Bayesian additive regression trees are similar to boosting in that decision trees are being successively estimated, this time in the form of particle Gibbs sampling of decision trees, with priors set over tree parameters (splitting structure, predictor selection, tree depth, residual variance) as regularization and Bayesian backfitting techniques to draw new trees based on the residual. This semi-parametric approach uses Markov Chain Monte Carlo procedures to draw additive trees as the fixed effects model components (*dbarts* ^29^), with subsequent Hamiltonian Monte Carlo (HMC) draws via Stan, to update patient/batch-level random intercepts ^30^.
4. **Bayesian generalized linear mixed-effects model (BGLMM) with Horseshoe Prior**: Implementation of Bayesian logistic regression using the *brms* package ^31^, where model parameters/random intercepts are drawn using HMC, and horseshoe shrinkage prior with documented performance over Lasso priors to effectively remove irrelevant/collinear genes ^32^.
5. **Regularized BGLMM with Interactions (BGLMM-Int):** A hybrid approach between mixed-effects CART methods and generalized linear procedures utilizes SHAP to extract the top 10 salient interactions from the GPBoost model and adds proposed interactions (robust to clustering; **Figure 1E**) as terms in the BGLMM Horseshoe model, under the premise that these interactions help the GLMM recover any performance gap for their MEML counterparts ^33^.
6. **BGLMM with Gaussian Process (BGLMM-GP):** Similar to GPBoost-Coords, replacing the gradient boosting decision trees with a linear functional: *logit*(*p*_*i*_) = β*x*_*i*_ + *b*^(1)^_[*i*]_ + *b*^(2)^_[*i*]_.
7. **Random Forest (RF) and Extreme Gradient Boosting (XGBoost):** Fixed effects counterparts (FEML) to aforementioned MEML methods, fit using *scikit-learn* and *xgboost* packages ^28,34^.

To assess the predictive performance of each modeling approach, we performed ten-fold cross-validation and bootstrapped performance statistics (C-Statistic and Log-Loss) through iterative resampling of the held-out validation sets of each of the cross-validation folds before averaging. We focused on sensible/coarse selection of hyperparameters. Salient predictors for the CART methods were extracted using SHAP to estimate the local importance of each predictor and summed to form global attributions ^35^. Interaction effects were extracted using SHAP’s *TreeExplainer*^20^. After selecting predictors via Horseshoe Lasso variable selection and standardizing, posterior draws of unpenalized BGLMM model parameters were used to assess and visualize significant predictors.

## 4. Results

### 4.1. Macro: Inter-Tumoral Prediction

The ability to characterize ROI out-of-batch/sample (*OOS*) indicates the potential to assess new cases prospectively for macro-architectural characteristics. Here, we expected within-sample (*WS*) performance to exceed predictions made on held-out patients. As expected, the prediction of whether an immune cell ROI was embedded in the interface with tumor exhibited greater performance within-sample across all approaches. MEML methods, taken as a group, slightly outperformed fixed effects approaches for macro-architectural prediction (**Table 1**). Surprisingly, generalized linear mixed effects approaches outperformed both the fixed effects and MEML methods, with slightly greater performance assigned to the BGLMM model, which utilized interactions detected from the GPBoost model (**Table 1)**. Adding spatial dependence to the model in the form of Gaussian Processes did not improve predictive performance for GPBoost and BGLMM.

**Table 1:** Bootstrapped model performance comparison, using C-Statistic/Area Under the Curve (AUC) and Log-Loss

|  |  | Fixed Effects |  | MEML |  |  | Bayesian Generalized Linear Mixed Models |  |  |
| --- | --- | --- | --- | --- | --- | --- | --- | --- | --- |
| Task | AUC ± SE | RF | XGBoost | GPBoost | GPBoost-Coords | SP-BART | BGLMM | BGLMM-Int | BGLMM-GP |
| Macro | OOS | 0.759±0.01 | 0.747±0.01 | 0.752±0.01 | 0.747±0.01 | 0.772±0.01 | 0.778±0.01 | **0.785±0.009** | 0.777±0.01 |
| | WS | 0.781±0.009 | 0.773±0.01 | 0.782±0.009 | 0.78±0.009 | 0.784±0.009 | 0.788±0.009 | **0.802±0.009** | 0.786±0.01 |
| METS | Overall | 0.909±0.006 | 0.951±0.004 | **0.971±0.003** | n/a | 0.897±0.006 | 0.852±0.008 | 0.896±0.006 | n/a |
| | Intra | 0.834±0.014 | 0.849±0.014 | **0.89±0.011** | n/a | 0.867±0.012 | 0.848±0.013 | 0.877±0.012 | n/a |
| | Inter | 0.881±0.013 | 0.866±0.013 | **0.899±0.012** | n/a | 0.874±0.012 | 0.836±0.014 | 0.881±0.012 | n/a |
| | Away | 0.827±0.014 | 0.842±0.014 | **0.895±0.011** | n/a | 0.887±0.011 | 0.885±0.012 | 0.885±0.011 | n/a |
| Task | Log-Loss ± SE | RF | XGBoost | GPBoost | GPBoost-Coords | SP-BART | BGLMM | BGLMM-Int | BGLMM-GP |
| Macro | OOS | 0.546±0.008 | 0.747±0.025 | 0.572±0.013 | 0.578±0.014 | 0.547±0.013 | 0.535±0.01 | **0.528±0.011** | 0.536±0.01 |
| | WS | 0.542±0.017 | 0.701±0.025 | 0.535±0.013 | 0.538±0.013 | 0.526±0.011 | 0.522±0.01 | **0.508±0.01** | 0.524±0.01 |
| METS | Overall | 0.485±0.005 | 0.284±0.013 | **0.248±0.007** | n/a | 0.409±0.014 | 0.455±0.01 | 0.393±0.01 | n/a |
| | Intra | 0.544±0.009 | 0.509±0.027 | **0.422±0.014** | n/a | 0.454±0.021 | 0.479±0.017 | 0.444±0.017 | n/a |
| | Inter | 0.507±0.009 | 0.499±0.032 | **0.412±0.017** | n/a | 0.457±0.026 | 0.484±0.017 | 0.429±0.019 | n/a |
| | Away | 0.559±0.01 | 0.559±0.031 | **0.428±0.017** | n/a | 0.442±0.023 | 0.441±0.015 | 0.434±0.015 | n/a |

Top predictors for tumor interface macro-architecture between the GPBoost model and BGLMM model were concordant (e.g., PanCk. CD127, CTLA4, CD44) and of similar directionality (**Figure 2A,C**). IgG isotope technical factor Ms-IgG2A was found to be in the top half of predictor for GPBoost, though effects were largely negligible and undetected by the BGLMM. Some of the interactions extracted from the GPBoost model were found to be some of the top predictors for the BGLMM-Int model (e.g., PanCk:CD68, PanCk:CD34) (**Figure 2B-C; Table 2)**, indicating the relevance of MEML extracted interactions in capturing macro-architectural associations.

**Figure 2:**
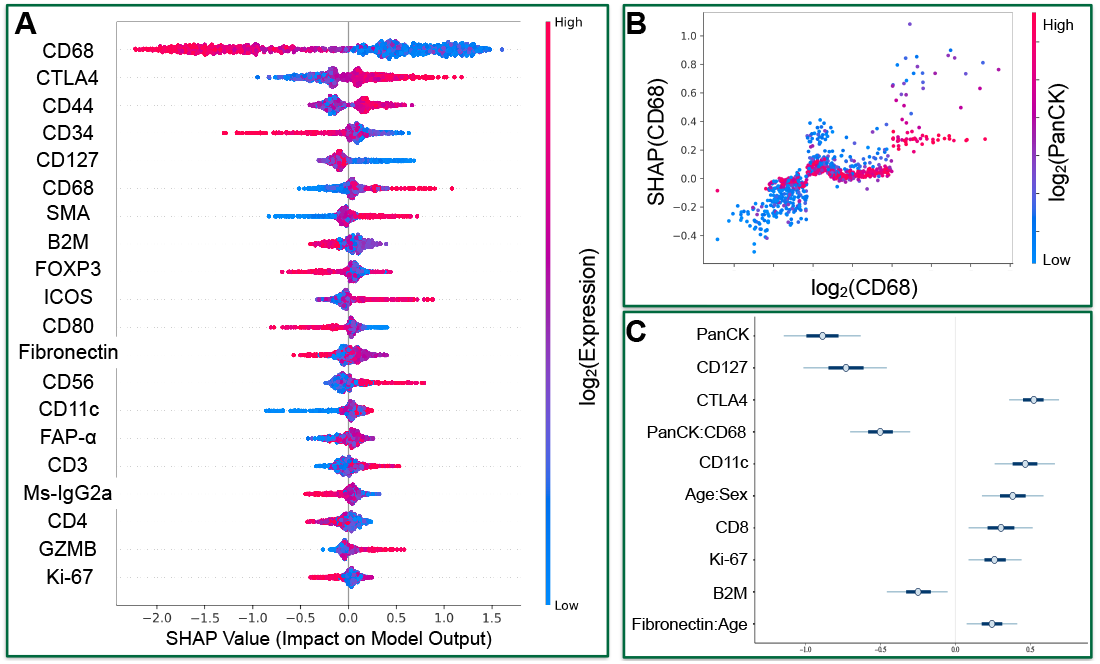
Macro Predictions: A) SHAP summary plot of top GPBoost terms; each point represents value of specific predictor for ROI; color indicates predictor’s value; positive SHAP value (x-axis) indicates how related to tumor interface; B) GPBoost SHAP partial dependency of CD68 interaction with PanCK; C) Posterior distributions of unpenalized BGLMM-Int predictors (thick and thin bars represent 50% and 90% credible intervals respectively)

**Table 2:** Top terms from BGLMM-Int models; GPBoost-extracted interactions are emphasized in bold

| Macro |  | Top 7 Ranked BGLMM-Int Terms |  |  |  |
| --- | --- | --- | --- | --- | --- |
| OOS | WS | METS |  |  |  |
|  |  | Overall | Intra | Inter | Away |
| PanCk | PanCk | **CD20:Age** | Beta-2-microglobulin (B2M) | **CD68:CTLA4** | CD25 |
| CTLA4 | CD127 | CD14 | CD45 | CD11c | CD44 |
| **CD68:PanCk** | CTLA4 | Age | CD8 | CTLA4 | FAP-α |
| CD34 | **PanCk:CD68** | CD68 | FOXP3 | CD14 | Fibronectin |
| **PanCk:CD27** | CD44 | CD11c | **B2M:FOXP3** | GZMB | CD68 |
| CD68 | **PanCk:CD34** | **Fibronectin:Age** | Age | CD68 | Age |
| CD27 | CD68 | Ki-67 | **CD8:CD45** | **CD14:CD66b** | **FAP-α:CD163** |
|  | CD56 | **Age:Sex** |  | Age | CD163 |
|  | CD34 | CD8 |  | **CD14:CTLA4** | PanCk |
|  | **PanCk:CD56** | Beta-2-microglobulin (B2M) |  | CD66b | **PanCk:Age** |
|  |  | CD20 |  | **GZMB:Age** |  |
|  |  | **B2M:Age** |  |  |  |
|  |  | Fibronectin |  |  |  |
|  |  | Sex |  |  |  |

### 4.2. METS: Nodal and Distant Metastasis Prediction

The greatest performance differences between fixed effects methods and MEML methods were noticed across all metastasis prediction tasks, where the GPBoost method obtained as little as a 2% boost in C-statistic to as much as a 9% performance gain versus RF and XGBoost. Within batch, it was easier to detect metastasis when all macro-architectural regions were utilized, while performance was roughly equivalent for each of the subregions (**Table 1**). Since performance of GPBoost exceeded that of all other approaches, the model may have learned significant interactions between genes that were invariant to batch differences. Using the detected interactions to supplement the BGLMM model allowed the BGLMM interaction model to vastly exceed the performance of its non-interaction counterpart and obtain performance competitive with GPBoost (**Table 1**). Across all tasks, the BGLMM model mostly outperformed its CART counterpart BART

GPBoost and BGLMM derived some concordant predictors (e.g., CD20, CD14), while others (e.g., FOXP3) were not found in the BGLMM model (**Figure 3A,C**). IgG isotope technical factor Ms-IgG1 was found to be in the top half of GPBoost predictors; effects were largely negligible and undetected by the BGLMM. The GPBoost model was also able to detect important interactions, (e.g., effect modification of markers CD20 and Fibronectin by age) (**Figure 3B-C; Table 2**) ^36^.

**Figure 3:**
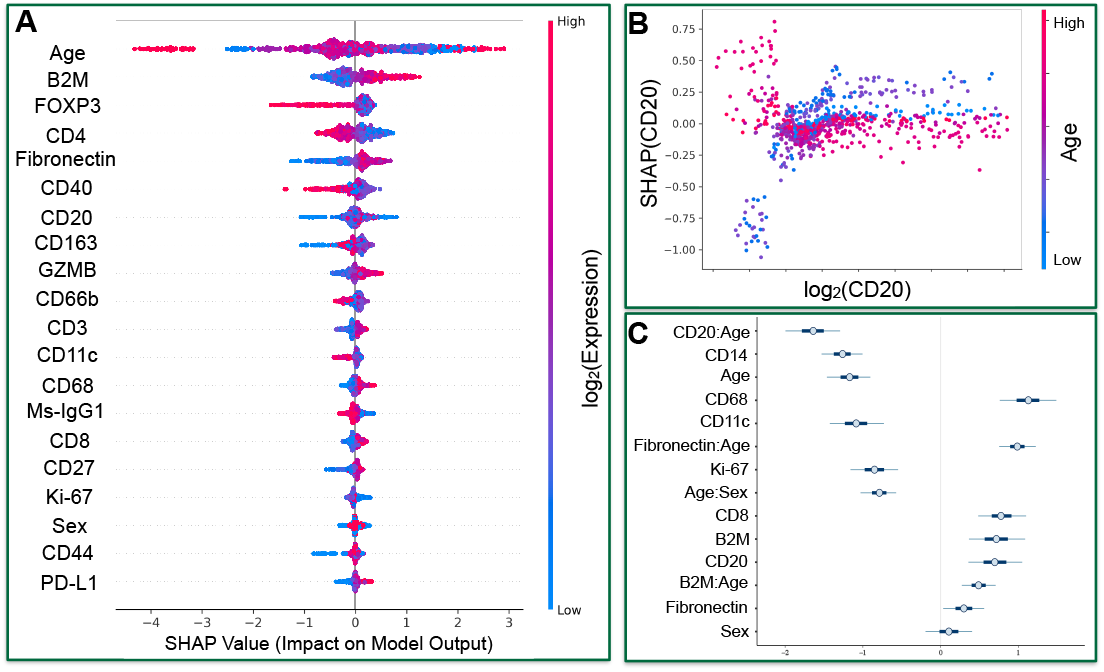
METS Predictions: A) SHAP summary plot of top GPBoost terms; each point represents value of specific predictor for ROI; color indicates predictor’s value; positive SHAP value (x-axis) indicates how related to tumor interface; B) GPBoost SHAP partial dependency of CD20 interaction with age; C) Posterior distributions of unpenalized BGLMM-Int predictors (thick and thin bars represent 50% and 90% credible intervals respectively)

## 5. Discussion

Spatial localization of gene and protein markers provides additional resolution on disease pathogenesis and etiology, which may soon lead to more informative and promising diagnostic and prognostic decision aids. While failing to account for repeated measurements in this setting may threaten to preclude classifier deployment, incorporating mixed-effects modeling into machine learning model development may derive potential additional clinically relevant disease predictors due to nonlinear interaction modeling while accounting for batch effects. In this study, we performed a preliminary assessment and comparison of different generalized linear and CART models for deployment as classifiers to assess macro-architectural characteristics and potential for undergoing nodal or distant metastasis. Results from our study indicate that classifier development for spatial omics technologies such as GeoMX DSP or Visium’s Spatial Transcriptomics may benefit by taking into account repeated measurements on the case and batch level. At minimum, mixed effects classification algorithms in spatial omics studies may detect complex gene-gene and gene-phenotype interactions that can be applied towards clinical findings (e.g., reportable risk estimates). For instance, the BGLMM-Int model indicated that the presence of CD20 (B-cells) was more predictive of metastatic potential in older individuals compared to younger individuals, while the presence of Fibronectin indicated the opposite effect (more predictive of metastasis in older individuals). We plan to further investigate the clinical utility of such identified interaction effects.

The added advantage of employing an MEML model is the ability to extract an interaction that the generalized linear approach can utilize, which may outperform generalized linear approaches without interactions, often unfairly reported in algorithm comparison studies. In this study, using this hybrid approach of informing interactions through MEML methods yielded BGLMM models that could outperform fixed effect and MEML methods due to simpler decision boundaries (line instead of stepwise function) and less biased sampling of the posterior of the parameters.

We acknowledge a few study limitations. Namely, we performed very coarse hyperparameter searches and other notable MEML methods ^27,29,37–40^ were omitted due to overlapping design. Additional algorithmic fine-tuning may demonstrate superior performance that is different from that reported in this work. The role of batch imbalance (association of batch with an outcome) has been well explored as it degrades the model performance in bulk expression datasets. However, its effects, particularly on intra-batch normalization and estimation of random effects, are unclear with the presented data and demand further exploration. We chose not to include batch effect correction using ComBat to limit the number of comparisons^41^. Regarding the batch effects, other than modeling Gaussian Processes nested within random intercepts, we did not include any random slopes in any of the MEML and BGLMM models, nor did we consider multiple hierarchy levels simultaneously (random intercept for batch and patient/biopsy). METS predictions were made within-batch and are not an accurate reflection of deployment on new samples. We did not incorporate time from initial biopsy to development of local or distant METS, though we plan to comment on the temporality of such associations in follow-up clinical findings. Projective predictive feature selection can help select more relevant BGLMM predictors ^42,43^. We neither performed nor compared findings from univariable analyses with multivariable analyses, which is outside the study scope. We also noticed that there was no gain in performance from using the ROI spatial coordinates, which may have been an artifact of implementational difficulties, the low sample size of ROIs and/or selection/learning of improper kernels. Spatial autocorrelation is often studied in spatial omics analyses, and, in the future, we expect to include an in-depth assessment of spatial effects.

In the future, we aim to refocus these findings for a more clinical setting after further algorithmic fine-tuning and orthogonally validation of our research results using immunohistochemistry (IHC). For prospective deployment of these technologies, we envision IHC serving as a low-cost alternative and potentially less encumbered by batch effects to rapidly spatially assess markers that have demonstrated utility for identification of colon metastasis. Such data modalities require standardization of collection/analysis processes (e.g., automated segmentation of macro-architecture (*intra, inter, away*) with deep learning algorithms, assessment of potential for bias from semi-subjective/automated selection of ROI) ^44^. Application of chemical reagents demands further adjustment to ensure meaningful deployment. Regarding further methods development, aggregation across macro-architectural regions may be explored using graph neural networks, which can pool spatial omics information across these regions to form a succinct vectorial representation of patient’s overall profile that can be integrated with H&E imaging modalities. Simulation studies may provide further validation of the utility of MEML methods for the assessment of spatial omics technologies.

## 6. Conclusion

In this work, we demonstrated the potential utility of MEML methods for the assessment of spatial protein markers collected using the Nanostring GeoMx DSP. We illustrated that MEML methods obtain superior within-batch performance for Colon metastasis prediction versus fixed effects and generalized linear methods. Furthermore, MEML methods, when combined with generalized linear modeling, may lead to clearer communication of significant spatial disease associations in clinical research studies through the extraction of key complex interactions. Further standardization of data collection, normalization, and analysis are potential opportunities to consider spatial omics machine learning technologies for meaningful clinical deployment.

## 7. Acknowledgements

We acknowledge James O’Malley, Robert Frost, Prajan Divakar, and Christian Haudenschild for leadership, support and discussion, and support of the Laboratory for Clinical Genomics and Advanced Technology in the Department of Pathology and Laboratory Medicine of Dartmouth Hitchcock Health System and Pathology Shared Resource, at the Norris Cotton Cancer Center at Dartmouth with NCI Cancer Center Support Grant 5P30 CA023108-37. Additional funding support from Burroughs Wellcome Fund Big Data in the Life Sciences training grant, Dartmouth College Neukom Institute for Computational Science CompX awards, and NIH grant R01CA216265.

*Preprint of an article published in Pacific Symposium on Biocomputing © 2021 World Scientific Publishing Co., Singapore, http://psb.stanford.edu/*

## Footnotes

† Code available on GitHub at the following URL: https://github.com/jlevy44/MEML_Colon_DSP_METS. Data available on reasonable request due to privacy/ethical restrictions.

